# Rethinking suicide Thi4 thiazole synthases: comparative genomic insights and pilot functional evidence

**DOI:** 10.1101/2025.09.23.678117

**Authors:** Edmar R. Oliveira-Filho, Kristen Van Gelder, David Obe, Cătălin Voiniciuc, Mark A. Wilson, Andrew D. Hanson

**Affiliations:** Horticultural Sciences Department, University of Florida, Gainesville, FL 32611, USA; Department of Biochemistry & Redox Biology Center, University of Nebraska, Lincoln, NE 68588, USA

**Keywords:** gene clusters, sulfur transfer chain, thiamin biosynthesis, DUF6775

## Abstract

Suicide thiazole synthases (Thi4) are mononuclear metal enzymes that form the thiazole moiety of thiamin from NAD^+^, glycine, and a sulfur atom that is stripped from an active-site cysteine residue, causing enzyme inactivation. Comparative genomic analysis indicates that prokaryotic Thi4 genes often cluster on the chromosomal regions encoding ThiS, ThiF, and other proteins that can produce, relay, or use persulfide or thiocarboxylate sulfur. This genomic evidence suggests that certain suicide Thi4s might use a persulfide or thiocarboxylate as sulfur donor instead of the active-site cysteine – i.e., that they can operate in a non-suicide mode – and that a metal cofactor reservoir supports Thi4 function. To explore these possibilities, we performed proof-of-concept experiments using *Escherichia coli* as a heterologous platform. A representative bacterial Thi4 that clustered with *thiS* and *thiF* complemented an *E. coli* Δ*thiG* (thiazole auxotroph) single mutant better than a Δ*thiG* Δ*thiF* Δ*thiS* triple mutant, consistent with predicted interactions with the host sulfide transfer chain. Collectively, this evidence indicates that suicide Thi4s may not necessarily operate suicidally and highlights genomic and structural clues that warrant deeper biochemical investigation.

## 1. Introduction

Plants, fungi, and certain prokaryotes produce the thiazole moiety of thiamin via a suicide thiazole synthase (Thi4; EC 2.4.2.60).^1-4^ Yeast and plant suicide Thi4s use an active-site cysteine (Cys) residue as sulfur donor for the reaction; whereby the cysteine is converted to dehydroalanine, inactivating the protein after a single reaction (**Figure 1a**).^2,5^ The presence of this active-site Cys distinguishes suicide Thi4s from catalytic Thi4s (EC 2.4.2.59), which have a histidine or other residue in place of Cys and use free sulfide as a sulfur donor.^6,7^ Yeast and plant suicide Thi4s are mononuclear metal enzymes, with Fe(II) as the most likely cofactor. Even though metalation of purified Thi4s with Fe(II) is required to detect biochemical activity, ^2,8^ the metal may be loosely bound because it could not be detected in two of the four reported suicide Thi4 crystal structures.

**Figure 1.**
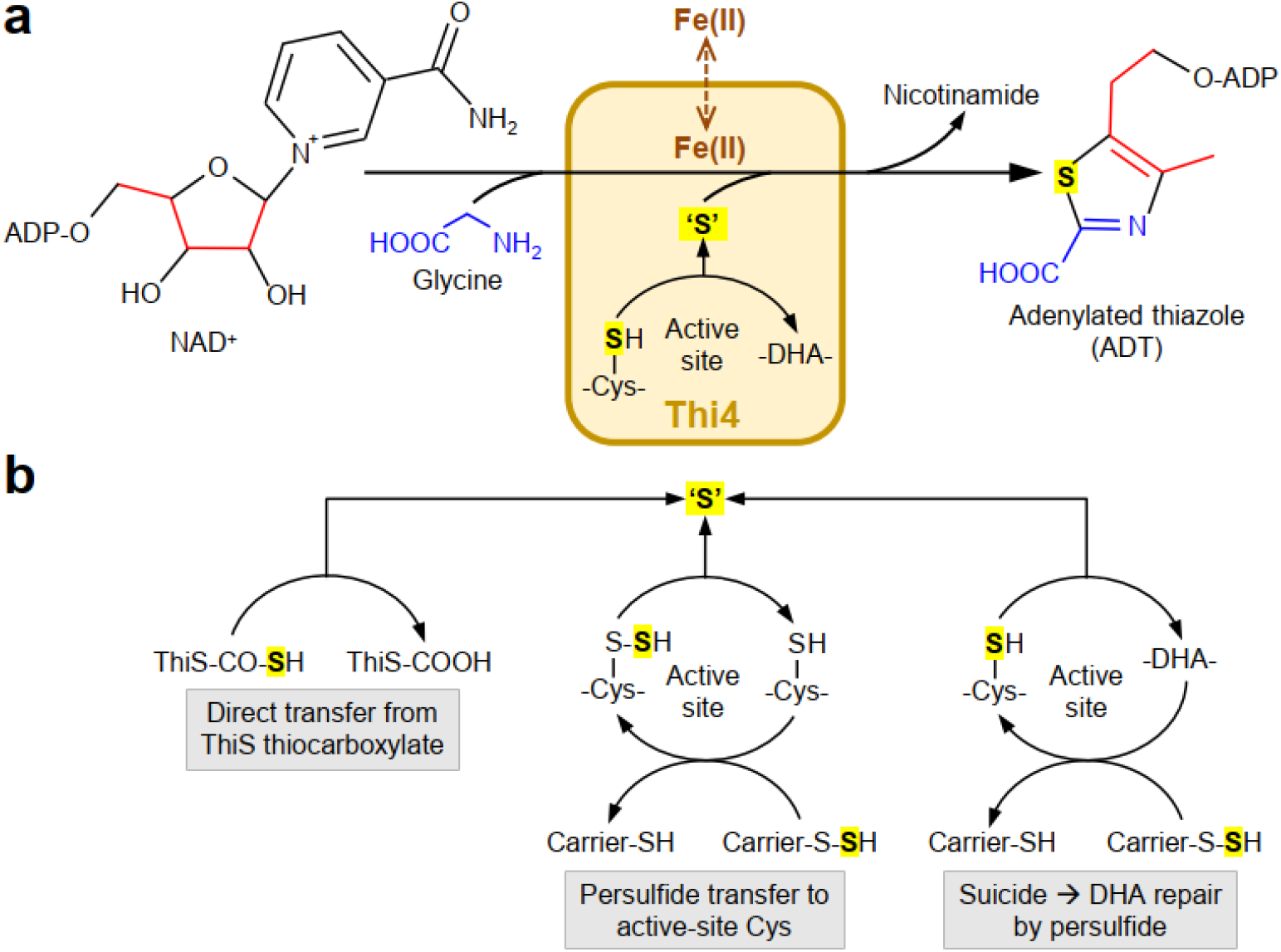
The complex reaction mediated by suicide Thi4s and hypothetical alternative sulfur donors for this reaction. (a) The known reaction. NAD+ and glycine are converted to the adenylated thiazole product (ADT) in an Fe(II) (or other metal)-dependent reaction in which the sulfur donor is an active-site Cys residue. This residue is converted to dehydroalanine (DHA). Red and blue colors show origins of the thiazole ring. The metal is weakly bound and may dissociate from the protein during the reaction. (b) Hypothetical alternative sulfur donors, based on comparative biochemistry.

Comparative biochemistry suggests that Thi4s with an active-site Cys could, in some cases, operate in a non-suicide mode, either by using an alternative sulfur donor that leaves the Cys intact or by repairing the dehydroalanine residue left after the reaction (**Figure 1b**). Precedents for these hypothetical options are as follows. The bacterial thiazole synthase (ThiG) (EC 2.8.1.10), which is unrelated to Thi4 and forms the thiazole product via a different reaction, provides a direct evidence for an alternative sulfur donor.^1^ The proximal sulfur donor for ThiG is a C-terminal thiocarboxylate group on the ThiS sulfur carrier protein ThiS (ThiS); this group is formed from a persulfide borne on the sulfur carrier protein ThiI (ThiI) in a reaction mediated by the adenylyltransferase (ThiF). The ultimate source of the persulfide sulfur, as in other sulfur-relay systems, is the action of a Cys desulfurase on free Cys.^1,9,10^ Another precedent is 2-thiouridine biosynthesis by MnmA, for which the proximal sulfur donor is a persulfide group on the sulfur carrier protein TusE.^9^ In these cases, genes encoding sulfur relay components often cluster on the chromosome with the gene for ThiG or MnmA, i.e. the enzyme that synthesizes the sulfur-containing product. A precedent for the repair of a dehydroalanine residue by a persulfide comes from the sacrificial sulfur insertase LarE from *Lactobacillus plantarum*; in this case the persulfide group is on a small-molecule thiol (Coenzyme-A), not on a protein carrier.^11^

Comparative biochemistry also suggests that suicide Thi4s, like other metalloenzymes, could be supported by metallochaperone proteins that deliver metal ions to their specific target and maintain metal homeostasis. Examples include the iron-trafficking chaperone HcdG, which delivers iron to the [Fe]-hydrogenase Hmd,^12^ and the nickel-binding protein HypA, which transfers a nickel ion to the catalytic core of the [NiFe]-hydrogenase.^13^ Like the sulfur-relay genes above, the genes encoding these chaperones are often clustered in operonic arrangements with the genes for the enzymes they service.^12,13^

These comparative biochemical and genomic considerations led us to briefly survey the genomic context of representative prokaryote suicide Thi4 genes. We saw obvious clustering with genes coding for various sulfur-relay proteins and their client biosynthetic enzymes, as well as with genes encoding for DUF6775, a domain of unknown function with predicted metal-binding residues. In this study, we applied the maxim^9^ that: “A genetic, genomic or proteomic…indication that…a protein homologous to another sulfur carrier protein has a role in a particular biological process should trigger a careful scrutiny for the involvement of a persulfide group in the biochemistry of the process”. We therefore combined comparative genomic analysis with pilot genetic and biochemical tests to explore the potential for alternative sulfur donors for prokaryotic Thi4 proteins with active-site Cys.

## 2. Materials and methods

### 2.1. Bioinformatics tools and databases

Prokaryote metagenomes and genomes were searched primarily in the Integrated Microbial Genomes & Microbiomes system (IMG/M) at the DOE Joint Genome Institute (https://img.jgi.doe.gov/m/)^14^ and secondarily in GenBank. The query was the suicide Thi4 protein sequence from Deltaproteobacteria bacterium RBG_16_54_11 (GenBank OGP80537); gene neighborhoods around similar suicide Thi4 genes were displayed in IGM/M via the ‘Show neighborhood regions with the same top COG hit (via top homolog)’ tool. Functional gene annotations were made or confirmed using the NCBI Conserved Domain Database (CDD).^15^

### 2.2. Plasmid constructs

All molecular manipulations followed standard protocols^16^ or reagent/kit manufacturers’ instructions. Codon-optimized nucleotide sequences with added compatible restriction sites were ordered from GenScript (Piscataway, NJ). Restriction cloning was used to clone genes using the following restriction enzymes (New England Biolabs, Ipswich, MA): *Eco*RI/*Xba*I for pBAD24; *Nde*I/*Hin*dIII for pET28b(+). Digestion products were gel-purified using GeneJET Gel Extraction Kits (Thermo Fisher Scientific, Waltham, MA), and assembled by T4 DNA Ligase (Thermo Fisher Scientific, Waltham, MA). Ligation products were transferred to *Escherichia coli* Top10 via electroporation using an *E. coli* Pulser apparatus (Bio-Rad Laboratories, Hercules, CA). Candidate clones (antibiotic-resistant) were then PCR-screened to select those with successful assembly. Oligonucleotide sequences are listed in **Table S1**. Recombinant plasmids were purified using GeneJET Plasmid Miniprep Kits (Thermo Fisher Scientific, Waltham, MA) and sequence-verified by Sanger or whole plasmid sequencing (Genewiz, South Plainfield, NJ).

### 2.3. Escherichia coli gene knockouts

*E. coli* MG1655 Δ*thiG* and Δ*thiF* Δ*thiS* Δ*thiG* mutants were generated through recombineering.^17^ For the Δ*thiG* single knockout, the deletion cassette was amplified by PCR from *E. coli* Keio JW5549-1 genomic DNA. To generate the Δ*thiF* Δ*thiS* Δ*thiG* mutant, a DNA fragment of the region encoding for the C-terminus of ThiE through to ThiF, ThiS, ThiG, and for the N-terminus of ThiH was amplified from *E. coli* MG1655 and cloned into pGE using NEBuilder HiFi DNA Assembly Cloning Kit (New England Biolabs, Ipswich, MA), generating pGE_*thiFthiSthiG*. A spectinomycin resistance cassette (Sm^R^) flanked by FRT sites was amplified from pTs and inserted into pGE_*thiFthiSthiG* using the same kit. The Sm^R^ cassette replaced *thiF, thiS*, and *thiG*, leaving 200-bp flanking sequences (homology arms) for *thiE* and *thiH* upstream and downstream of Sm^R^, respectively. Each deletion cassette was transferred via electroporation to *E. coli* MG1655 harboring pKD46. Recombinants were selected on LB (tryptone, 10 g/L; yeast extract, 5 g/L; and NaCl, 5 g/L) with kanamycin (50 µg/mL) or spectinomycin (100 µg/mL). Candidate clones were screened by colony PCR followed by agarose gel electrophoresis. pKD46 was purged from PCR-verified clones by overnight incubation at 42°C.

### 2.4. Functional complementation assays

*Methanobacterium* sp. MB1 (MB1) and *Candidatus Omnitrophica* bacterium isolate bin.255MB1 (cOb) THI4 encoding sequences were recoded for expression in *E. coli* (**Table S2**) and synthesized by GenScript (Piscataway, NJ). pBAD24 constructs harboring *thi4* were transformed into *E. coli* MG1655 Δ*thiG* and Δ*thiF* Δ*thiS* Δ*thiG*. pBAD24::Ta_*thi4* was used as a benchmark.^7^

To verify the production of soluble THI4 protein, recombinant MG1655 was cultured at 37°C and 250 rpm in 25 mL MOPS minimal medium^18^ supplemented with 0.2% glycerol, micronutrients,^19^ 100 µg/mL carbenicillin, and 100 nM thiamin. When OD_600_ reached ∼0.5, cultures were cold-shocked in an ice bath for 30 min, L-Arabinose was added to a final concentration of 0.02% (w/v), before an overnight incubation at 18°C, 180 rpm. Cells were harvested by centrifugation (5000*g*, 15 min, 4°C) and stored at -80°C. Cell pellets were sonicated in 100 mM potassium phosphate (pH 7.2), 2 mM β-mercaptoethanol buffer. Protein concentration was determined by Bradford assay, and protein profiles (10 µg per sample) were visualized by SDS-PAGE on 12.5% gels stained with GelCode Blue Safe Protein Stain (Thermo Fisher Scientific, Waltham, MA).

Functional complementation was evaluated on the MOPS minimal medium with the supplements described above. Single colonies were grown in 3 mL of medium containing 100 nM thiamin for 24 hours. These were sub-cultured twice on thiamin-free MOPS supplemented with 1 mM Cys, and 0.02 % (w/v) L-arabinose. Cells from the third culture were washed three times with basal MOPS (i.e., MOPS medium minus MgSO_4_, FeSO_4_, micronutrients, or thiamin) and used to prepare serial 10-fold dilutions starting with an OD_600_ of 1.5. These were spot-plated on MOPS agar plates with 1 mM Cys, 0.02% (w/v) L-arabinose, with or without 100 nM thiamin. Plates were incubated at 37°C for 3-4 days.

### 2.5. Recombinant DUF6775 production and bound metal analysis

Selected DUF6775-containing sequences were codon-optimized for *E. coli* (**Table S2**) and synthesized by GenScript (Piscataway, NJ). pET28b(+) constructs were transferred to *E. coli* BL21 (DE3) by electroporation. Recombinant BL21 was cultured on 50 mL LB plus kanamycin at 37°C, 250 rpm. When OD_600_ reached ∼0.5, cultures were cold-shocked in an ice bath for 30 min, induced with 100 µM IPTG, and incubated overnight at 18°C, 180 rpm. Cells were harvested by centrifugation (5000*g*, 15 min, 4°C) and stored at -80°C. Cell pellets were sonicated in 25 mM HEPES-NaOH (pH 7.5), 100 mM KCl, 0.5% Triton X-100, and 5 mM dithiothreitol. Non-soluble fractions were collected by centrifugation (25000g, 30 min, 4°C), washed three times with the same buffer with 1% Triton X-100, with a short sonication cycle (3x 10s on, 50s off) between washes (25000*g*, 15 min, 4°C); and one time with 25 mM HEPES-NaOH (pH 7.5), 1 M KCl, 5 mM dithiothreitol. The obtained pellet was then resuspended in 25 mM HEPES-NaOH (pH 7.5), 100 mM KCl, and 5 mM dithiothreitol. Protein profiles were visualized by SDS-PAGE on 12.5% gels stained with GelCode Blue Safe Protein Stain (Thermo Fisher Scientific). Protein concentration was adjusted to 1 mg/mL by measuring absorbance at 280 nM and calculating concentrations based on the peptide’s extinction coefficient (ProtParam).

Metals associated with recombinant DUF6775 recovered from inclusion bodies were measured using inductively coupled plasma-mass spectrometry (ICP-MS) using an Agilent 7500 cx ICP-MS equipped with a 96-well plate autosampler.^20^ Samples were lyophilized and then incubated overnight with 100 μL of 70% trace metal-grade nitric acid at 65°C. After acid digestion for 14 hours, the samples were cooled to room temperature, centrifuged, and diluted 10-fold with 2% nitric acid spiked with 50 μg/L gallium (Ga) as the internal ICP-MS standard. The final diluted sample was injected into the ICP-MS autosampler for analysis, following the method described by Malinouski et al.^21^ ICP-MS measurements were performed in mix reaction mode (3.5 mL H_2_ and 1.5 mL He per minute) to eliminate polyatomic interferences in the collision cell. The samples were loaded using an autosampler (ESI, Omaha, NE) and injected through a 6-port injection valve with a 125 μL sample loop at a flow rate of 55 μL/min. The following ICP-MS operating conditions were used: Ar carrier flow: 1.0 L/min; Ar make-up flow: 0.1–0.2 L/min; forward power: 1,500 W; Ar plasma gas: 15 L/min; Ar auxiliary gas: 1 L/min. Metal concentrations were determined using an external calibration curve, with Ga (50 ppb) as the internal standard.^22^ Element ratios were calculated as the ratio of the element concentration to the protein concentration. All solutions for metal analysis were prepared using metal-grade water and nitric acid.

### 2.6 Structural model prediction

The structure of the conserved hypothetical archaeal DUF6775 protein was predicted using the AlphaFold 3 server.^23^ The resulting model exhibited a high overall confidence, with a mean predicted Local Distance Difference Test (pLDDT) score of 94.6. Regions of lower confidence (70 > pLDDT > 50) were localized to helices α7 and α8, as well as loop regions corresponding to loops 8, 10, 15, and 17. Proteins with experimentally determined structures similar to the predicted structure of DUF6775 were identified using the DALI server.^24^ Two structural homologs were identified: the zinc-dependent peptidase from *Methanocorpusculum labreanum* (PDB: 3LMC, UniProt: A2SQK8) and Archaemetzincin from *Methanopyrus kandleri* (PDB: 2X7M),^25^ with DALI Z-scores of 13.9 and 14.3, respectively. These homologs were used as structural templates to infer the likely metal-binding site in DUF6775. The identity of the bound metal in 2X7M has been experimentally verified via anomalous difference Fourier maps,^25^ although the methods used to assign the metals in 3LMC are unknown, as there is no supporting publication.

All predicted models and structural comparisons were visualized and analyzed using UCSF ChimeraX version 1.9.^26^

## 3. Results and discussion

### 3.1. Neighboring-gene analyses of suicide Thi4s

The suicide Thi4 gene from *Deltaproteobacteria* bacterium RBG_16_54_11 is typical of those seen in our initial survey to cluster with sulfur-relay and DUF6775 genes (**Figure 2a**). This Thi4 gene was used as a query to extract gene neighborhood data from the IMG/M system^14^ and GenBank for >400 suicide Thi4 genes from bacteria and archaea belonging to at least 26 different phyla; **Figure 2a** shows representative cases. From these data, we identified genes that cluster consistently in the ten-gene window around the Thi4 gene (**Figure 2b** and **Table S3**).

**Figure 2.**
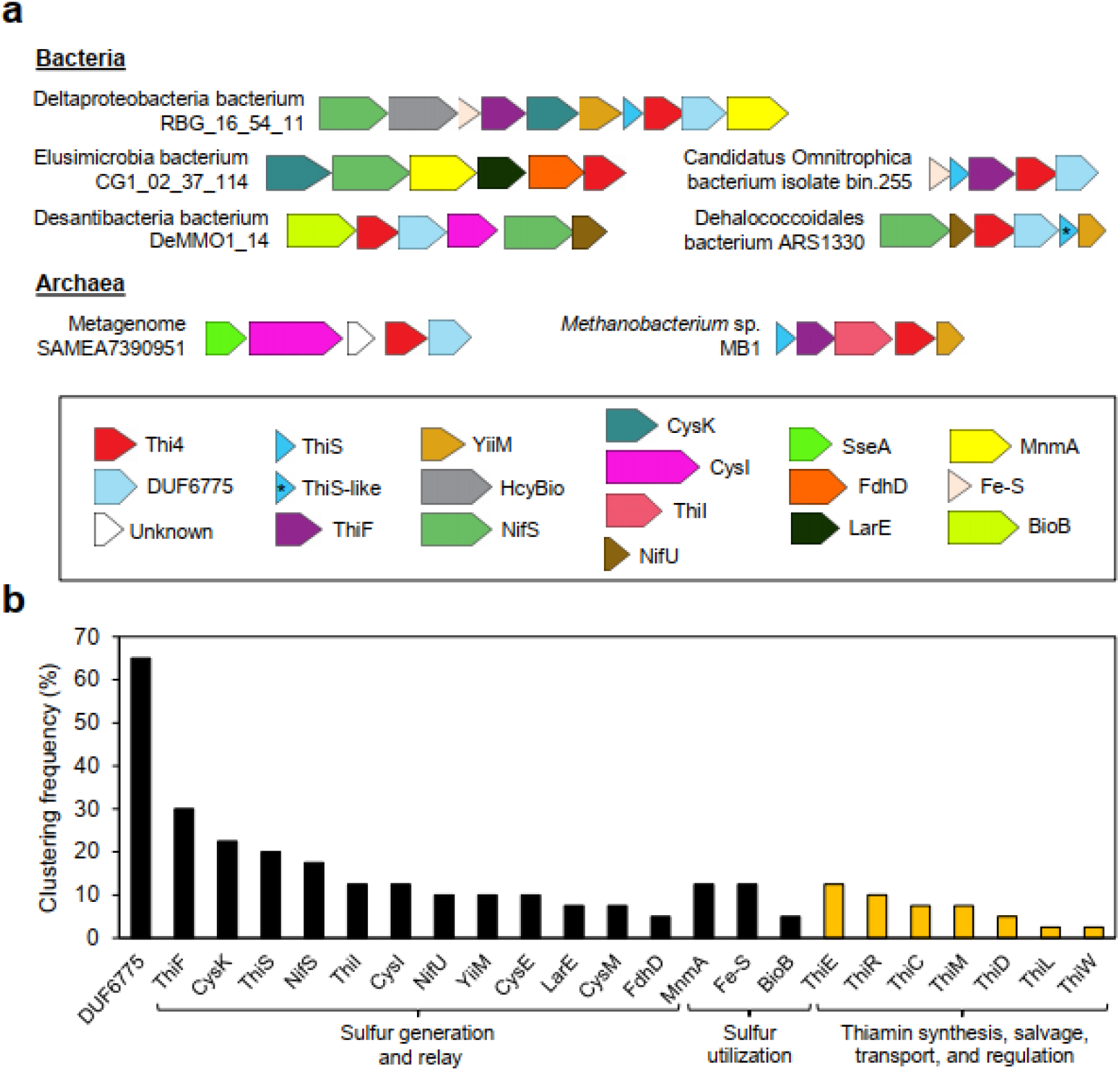
Gene neighborhoods around suicide Thi4s. (a) Representative examples from bacteria and archaea. (b) Frequencies (percent of metagenomes and genomes surveyed) with which genes for DUF6775, sulfur-generating enzymes, sulfur-relay proteins, sulfur-utilizing enzymes, and thiamin synthesis and salvage enzymes (ThiC, ThiD, ThiE, ThiL, ThiM), the carrier ThiW, and the regulator ThiR recur in the 10-gene window around Thi4. Abbreviations: ThiF, adenylyltransferase catalyzing adenylylation of the ThiS C terminus; CysK, Cys synthase; ThiS, sulfur carrier; NifS, Cys desulfurase; ThiI, sulfurtransferase; CysI, sulfite reductase; NifU, FeS cluster assembly scaffold protein; YiiM, MOSC (MOCO sulfurase C-terminal) domain protein; CysE, serine O-acetyltransfer-ase; LarE, sacrificial sulfur insertase; CysM, S-sulfocysteine synthase; FdhD, sulfurtransferase; MnmA, tRNA U34 2-thiouridine synthase; Fe-S, iron-sulfur cluster protein; BioB, biotin synthase; HcyBio, L-aspartate semialdehyde sulfurtransferase; SseA, 3-mercaptopyruvate sulfurtransferase.

Genes encoding DUF6775 were the most frequently clustered with Thi4, followed by a full range of genes for sulfur generation and relay proteins including ThiF and ThiS, and by genes for proteins that use the sulfur delivered by relays (MnmA, BioB, and various Fe-S proteins) (**Figure 2b**). These genes all clustered with Thi4 at similar or higher frequencies than thiamin synthesis, salvage, transport, and regulation genes (ThiC, ThiD, ThiE, ThiL, ThiM, ThiW, and ThiR) (**Figure 2b**). None of the genomes from **Figure 2a** encoded ThiG (**Table S4**), indicating that the Thi4-clustered ThiS and ThiF proteins in these genomes cannot be servicing ThiG. These patterns indicate possible functional relationships between suicide Thi4s and sulfur transfer components at least as strong as those with other thiamin-related enzymes, carriers, and regulators.^27^ The alternative mechanism used by the non-suicide Thi4 (sulfur donor or Thi4, see **Figure 1b**) cannot be distinguished by the comparative genomic data. Direct sulfur donation by ThiS-thiocarboxylate, as established for ThiG in thiamin biosynthesis,^28^ and persulfide-mediated repair of dehydroalanine, analogous to pathways in tRNA thiolation^29^ remain plausible. Given the *a priori* plausibility of non-suicidal Thi4 action and of the dependence of Thi4s on a metal chaperone (see Introduction), the above comparative genomic evidence led us to run pilot experiments to test both possibilities.

### 3.2. Complementation tests for interaction of Thi4s with E. coli sulfur carrier protein ThiS

A straightforward possibility suggested by the clustering of *Thi4* genes with *ThiS* and *ThiF* genes (**Figure 2**) is that, like ThiG, Thi4 can accept sulfur from the ThiS thiocarboxylate group formed by ThiF.^1^ We therefore sought Thi4s that are (i) encoded by genes clustered with *thiS* and *thiF* genes, (ii) from mesophiles (and hence likely to function in *E. coli*), and (iii) that express well as soluble proteins in *E. coli*. Thi4s from *Methanobacterium* sp. MB1 (MB1) and *Candidatus* Omnitrophica bacterium (cOb) isolate bin.255 met these criteria (**Figure 2a** and **Figure S1**). These two Thi4s were then tested for their ability to complement an *E. coli* single Δ*thiG* mutant or a triple Δ*thiG* Δ*thiF* Δ*thiS* deletion mutant. We generated these mutant combinations to investigate whether the selected Thi4 proteins require sulfur from the alien *E. coli* ThiS, which would result in stronger complementation in the single mutant than in the triple mutant. The catalytic *Thermovibrio ammonificans* TaThi4 was included as a benchmark because it uses free sulfide as sulfur donor^7^ and is consequently predicted to perform equally well in both mutant backgrounds.

The TaThi4 control worked as expected, and we found stronger complementation of the single Δ*thiG* mutant for cObThi4 but not for MB1Thi4 (**Figure 3**). Thus, as cells were serially passaged without thiamin, the growth of cells expressing cOb4Thi4 became significantly weaker in the triple mutant relative to the single mutant, which was not the case for MB1Thi4 or TaThi4 (**Figure 3a**). Weaker complementation by cObThi4 was also visible on agar plates using thiamin-depleted cells used as inoculum (**Figure 3b**). Because the native ThiS proteins of MB1 and cOb are similar to *E. coli* ThiS only in the C-terminal quarter (**Figure S2**), and ThiS family sulfur carriers can be specific to particular sulfur acceptor enzymes,^30^ lack of stronger complementation in the single mutant might arise from failure of MB1Thi4 to interact functionally with *E. coli* ThiS. Therefore, the negative result with MB1Thi4 does not necessarily exclude the ThiS of MB1 as a physiological sulfur donor for MB1Thi4 in the native host.

**Figure 3.**
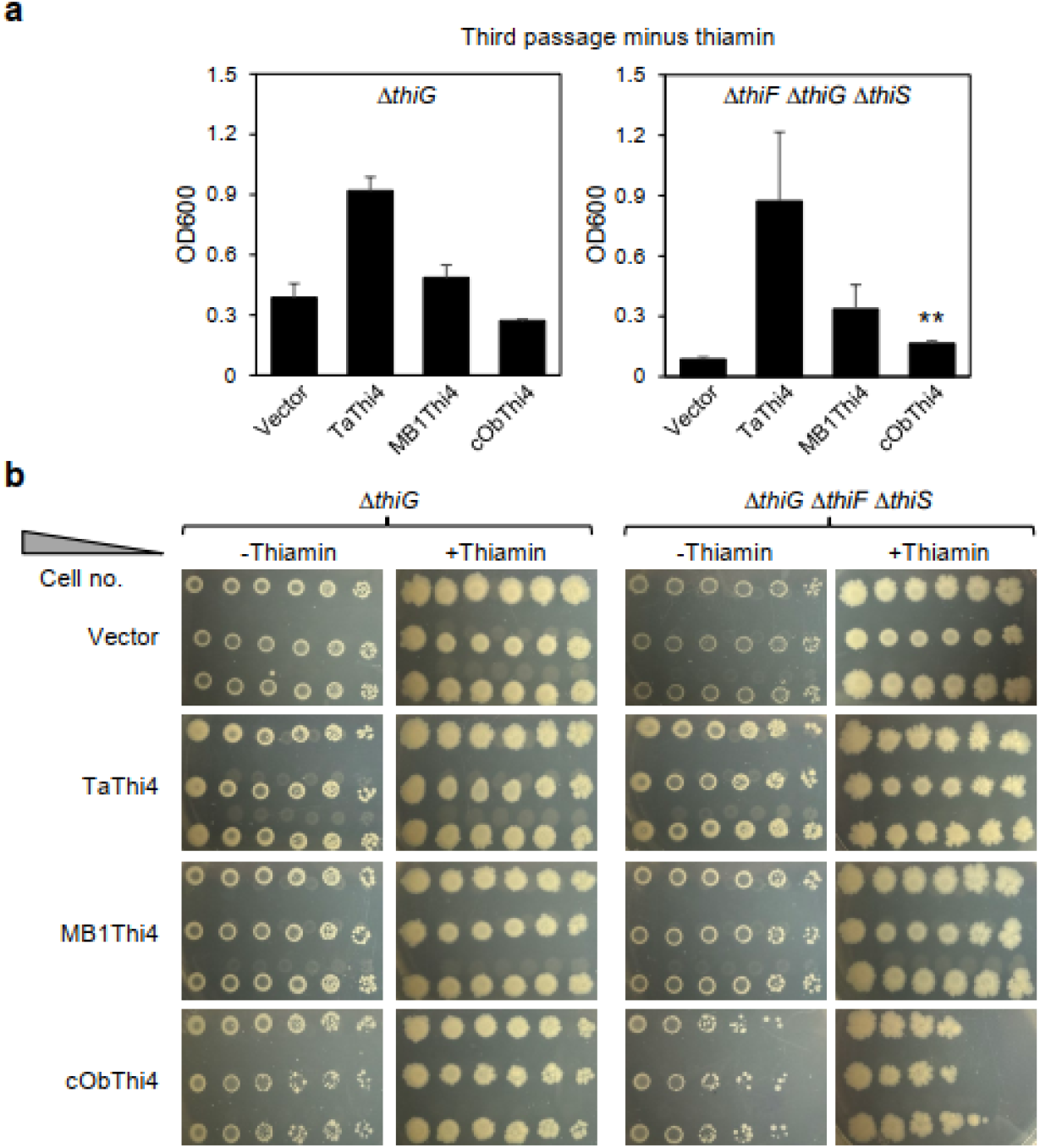
Complementation tests of interaction between two suicide Thi4s and the heterologous ThiF-ThiS system of *E. coli*. The *E. coli* host strains were deleted for *thiG* alone or for *thiF* and *thiS* as well as *thiG*. (a) Growth at 24 h of the liquid cultures minus thiamin used to inoculate the agar plate complementation tests. Error bars are SEM of three biological replicates. (b) Growth at 4 days on the complementation plates. Abbreviations: Vector, vector alone; TaThi4, non-suicide Thi4 from *Thermovibrio ammonificans* (benchmark); MB1Thi4, suicide Thi4 from *Methanobacterium* sp. MB1; cObThi4, suicide Thi4 from *Candidatus Omnitrophica* bacterium isolate bin.255. Note the poorer growth of cells expressing cObThi4 in the triple mutant host compared to the single mutant host in liquid medium minus thiamin (**, P<0.01) and on plates minus or plus thiamin, and the lack of such a difference for single and triple mutant cells expressing TaThi4 or MB1Thi4. Similar results were obtained in three independent experiments.

### 3.3 Predicted structure and pilot metal-binding tests of a DUF6775 protein

We also found that poorly characterized DUF6775 genes (no publications in PubMed or Google Scholar as of November 2025) cluster with Thi4s in two-thirds of the genomes analyzed (**Figure 2b**) and are annotated as putative metallopeptidases. Representative DUF6775 proteins have conserved Cys and histidine residues for potential metal binding (**Figure S3**), so we analyzed their structure with AlphaFold 3. The proteins chosen for analysis came from a mesophile archaeon (metagenome SAMEA7390951, SAMEA). The AlphaFold DUF6775 model has a three-layer αβα sandwich fold (**Figure 4**) characteristic of metallopeptidases within the collagenase catalytic domain superfamily (CATH 3.40.390.10). Related proteins with known structures include PDB 3LMC (Northeast Structural Genomics Consortium Target MuR16) and 2X7M,^25^ which contain bound metals (**Figure 4**). We note that the assignment of one of the metal sites in 3LMC as Fe is surprising given its 3xCys-1xHis tetrahedral coordination geometry and the assignment of Zn to the corresponding site of 2X7M. Unlike canonical archaemetzincins, which contain a HEXXHXXGX_3_CX_4_CXMX_17_CXXC zinc-binding motif comprising four strictly conserved Cys residues,^25^ DUF6775 has a distinct predicted metal binding site with three conserved Cys and one histidine (**Figure 4**), and it lacks the classical HEXXHXXGXXH/D motif that coordinates the second Zn^2+^ ion. These differences suggest a metal coordination environment in DUF6775 that is different from known zinc metallopeptidases with related structures.

**Figure 4.**
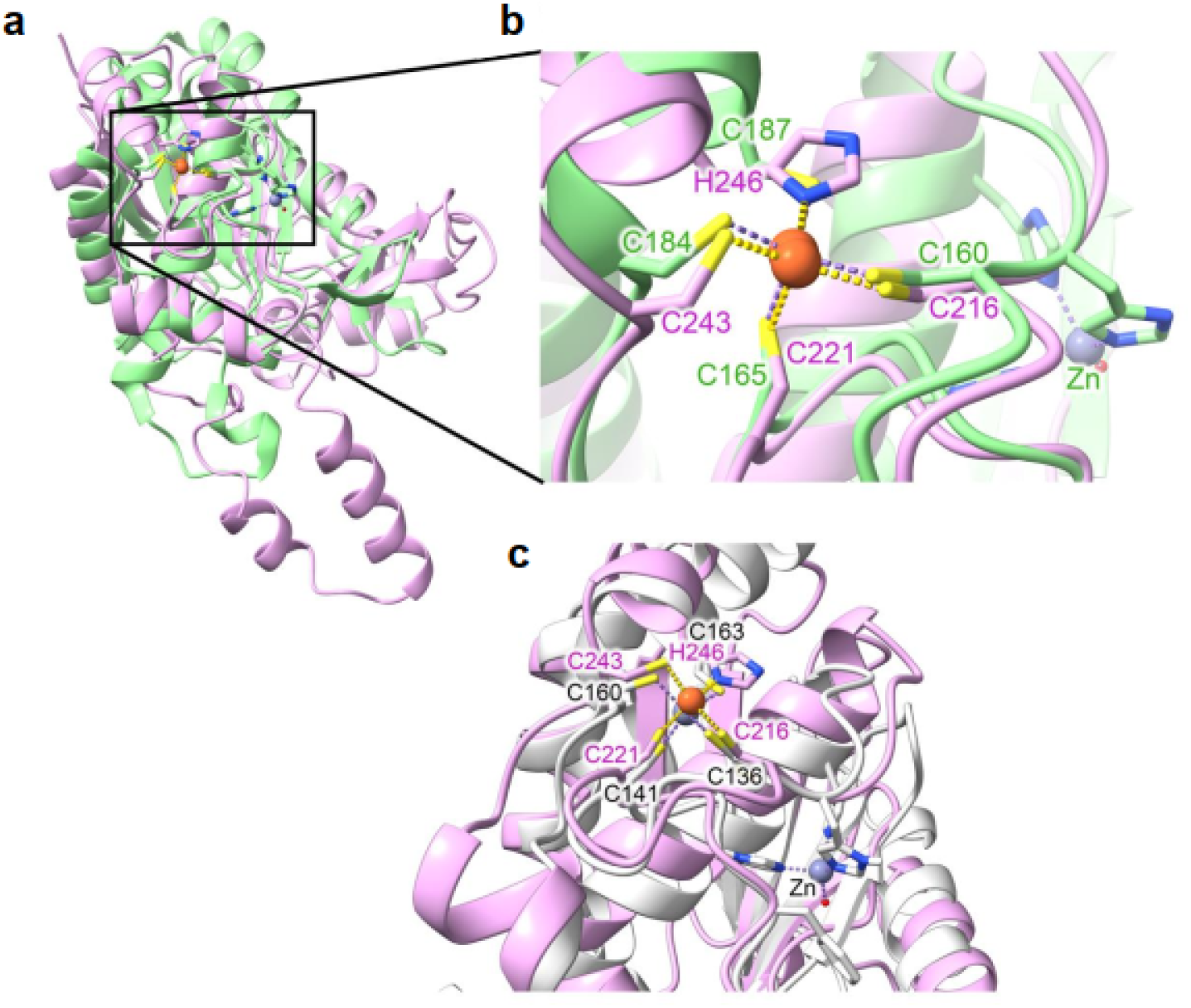
Predicted structure of DUF6775 and comparison to experimentally determined metallopeptidases. (a) Superposition of the AlphaFold3-predicted structure of DUF6775 (pink) and PDB 3LMC (green). 3LMC contains two metals: Fe (orange) and Zn (grey), although the Fe site is unusual and requires validation. (b) The coordination environment in a close-up inset. Note that the His residues that coordinate the second Zn in 3LMC are absent in DUF6775. (c) Superposition of DUF6775 (pink) and PDB 2X7M, the structure of archaemetzincin (AmzA) from *Methanopyrus kandleri*.^26^ The predicted metal in DUF6775 is orange, and the experimentally determined Zn in 2X7M are grey.

To experimentally investigate its ability to bind metals, a DUF6775 protein from the archaeal metagenome SAMEA7390951 was expressed in *E. coli*. The recombinant protein (∼33 kDa) was only found in the insoluble fraction of *E. coli* lysates, and was considerably enriched following an inclusion body preparation protocol (**Figure S4**). However, analysis of the inclusion body samples via inductively coupled plasma mass spectrometry (ICP-MS) did not identify any bound metals relative to the basal buffer control (**Table S5**). Expression trials were conducted with additional DUF6775 sequences obtained from diverse genomes and metagenomes, including *Dehalococcoidales* bacterium, *Hadesarchaea* archaeon DG-33-1 (2656880379), *Pyrococcus* sp. NA2 (650831530), and candidate division TA06 bacterium SM23_40. However, none yielded soluble protein (**Figure S5**). Future work should focus on optimizing the expression and purification of soluble, active recombinant DUF6775 proteins to identify the associated metal(s), to further investigate their function, and to test or refine the hypotheses presented here.

## 4. Conclusion

This work presents comparative genomic evidence indicating a possibility that, as already known for ThiG thiazole synthase,^6,7^ at least some suicide Thi4s can use a sulfur relay system to provide the sulfur atom needed for the thiazole ring, and thus are not necessarily doomed to sudden death. Pilot experimental data consistent with this possibility are also presented. Future biochemical studies that one of these Thi4s produces thiazole without consuming the active-site Cys will be needed to thoroughly test this hypothesis. The kinetic competitiveness between intramolecular sulfur donation from the active-site Cys and potential intermolecular sulfur transfer from ThiS, as well as the thermodynamic driving force for such reactions, remains unresolved. Future work should involve purified Thi4 and ThiS proteins under anaerobic conditions, coupled with mass spectrometry of reaction intermediates and active-site residues, to quantify reaction rates and determine whether sulfur relay can outcompete the canonical suicide pathway. While not definitive, these data warrant future, deeper investigation of the role of ThiS family proteins as potential sulfur donors for Thi4s, e.g. by constructing and expressing in *E. coli* synthetic operons containing a suicide Thi4 and the ThiS, ThiF, and other sulfur relay genes with which the Thi4 gene clusters in its native host (see **Figure 2a**).

Here, we also unveiled a close genomic association between suicide Thi4s and DUF6775 family proteins, which remain uncharacterized but feature similar motifs to metal-dependent peptidases. Although we could not produce soluble forms of DUF6775 in bacteria when using four diverse coding sequences, pilot ICP-MS analysis was conducted with samples obtained from inclusion bodies, from which we could not detect any bound metal. Recombinant proteins in inclusion bodies may be poorly folded or otherwise incapable of binding metal, so the lack of ICP-MS-detectable metal in our samples does not exclude the possibility that DUF6775 acts as a physiological cofactor for Thi4. It is worth noting that non-suicide Thi4s cluster strongly with genes encoding TRASH proteins known to bind various metals.^7^ Therefore, these exploratory data warrant further biochemical and genetic investigation to elucidate the functions of prokaryotic Thi4s and of their neighboring genes that encode uncharacterized domains.

## Author contributions

**Edmar R. Oliveira-Filho:** Conceptualization, Methodology, Investigation, Visualization, Writing – original draft. **Kristen Van Gelder:** Methodology, Investigation, Writing – review & editing. **David Obe:** Methodology, Investigation, Writing – review & editing. **Cătălin Voiniciuc**: Writing – review & editing. **Mark A. Wilson:** Conceptualization, Methodology, Investigation, Visualization, Writing – review & editing, Funding acquisition. **Andrew D. Hanson:** Conceptualization, Writing – original draft, Funding acquisition, Project administration.

## Supporting Information

**Figure S1**. SDS-PAGE analysis of the soluble fraction of Δ*thiG* and Δ*thiF* Δ*thiG* Δ*thiS E. coli* strains expressing selected *thi4*.

**Figure S2**. Alignment of *E. coli* ThiS with representative ThiS family or ThiS-like proteins from Figure 2a whose genes cluster with suicide Thi4 genes.

**Figure S3**. Alignment of representative DUF6775 proteins from **Figure 2a** whose genes cluster with suicide *thi4* genes.

**Figure S4**. SDS-PAGE analysis of recombinant DUF6775 protein from archaeal metagenome SAMEA7390951.

**Figure S5**. SDS-PAGE analysis of recombinant DUF6775 protein from different (meta)genomes.

**Table S1**. Primer sequences used in this work.

**Table S2**. Recoded nucleotide sequences of Thi4 and DUF6775 proteins tested in this study.

**Table S3**. Sulfide-related, thiamin-related, and DUF6775 genes in the 10-gene window around the suicide Thi4 gene in each of 40 genomes representative of the >400 genomes analyzed.

**Table S4**. Occurrence of Genes Coding for ThiG, ThiS, and ThiF Family Proteins in the Seven Representative Genomes from Figure 2a.

**Table S5**. Inductively Coupled Plasma–Mass Spectrometry (ICP-MS) Analysis of DUF6775 Protein Samples.

## Acknowledgments

*We dedicate this work to the memory of Andrew D. Hanson, whose leadership, wisdom, and generosity enriched the lives of all who worked with him*.

We thank Dr. Javier Seravalli of the University of Nebraska Spectroscopy and Biophysics Core Facility for performing the ICP-MS measurements.

