## Supplemental Figure for "Rethinking suicide Thi4 thiazole synthases: comparative genomic insights and pilot functional evidence"

∆*thiG*

V	Ta	MB1	cOb

26

34

∆*thiF* ∆*thiG* ∆*thiS*

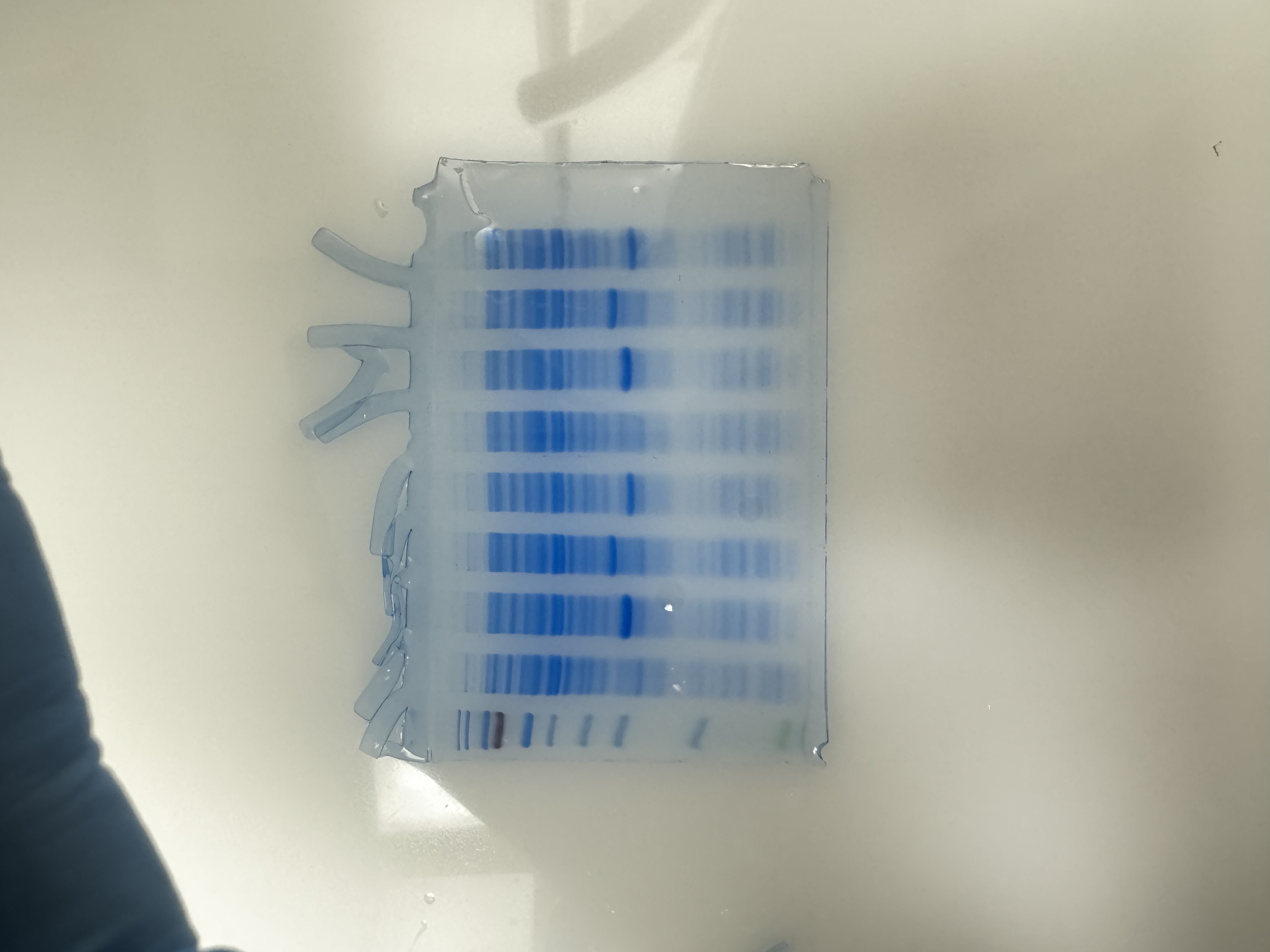

V	Ta	MB1	cOb

kDa

**Figure S1.** SDS-PAGE analysis of the soluble fraction of Δ*thiG* and ∆*thiF* ∆*thiG* ∆*thiS. E. coli* strains expressing *thi4* from *Thermovibrio ammonificans* (Ta), as a benchmark, shown to express well in *E. coli*,^7^ *Methanobacterium* sp. MB1 (MB1), or *Candidatus* Omnitrophica bacterium isolate bin.255 (cOb), or harboring vector alone (V). The gel was stained with GelCode Blue Safe Protein Stain.

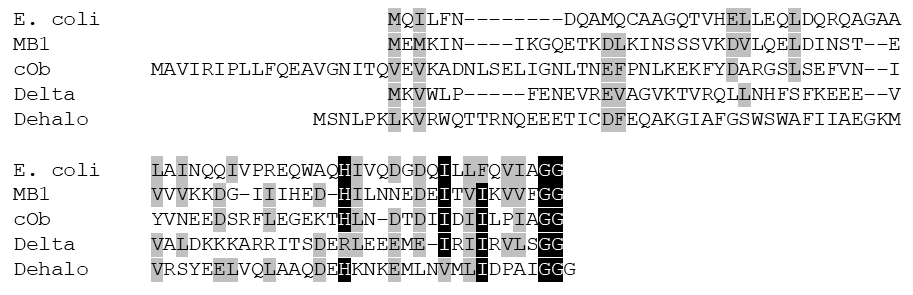

**Figure S2.** Alignment of *E. coli* ThiS with representative ThiS family or ThiS-like proteins from Figure 2a whose genes cluster with suicide *thi4* genes. Abbreviations: MB1, *Methanobacterium* sp. MB1; cOb, *Candidatus* Omnitrophica bacterium isolate bin.255; Delta, Delta-proteobacteria bacterium RBG_16_54_11; Dehalo, Dehalococcoidales bacterium ARS1330.

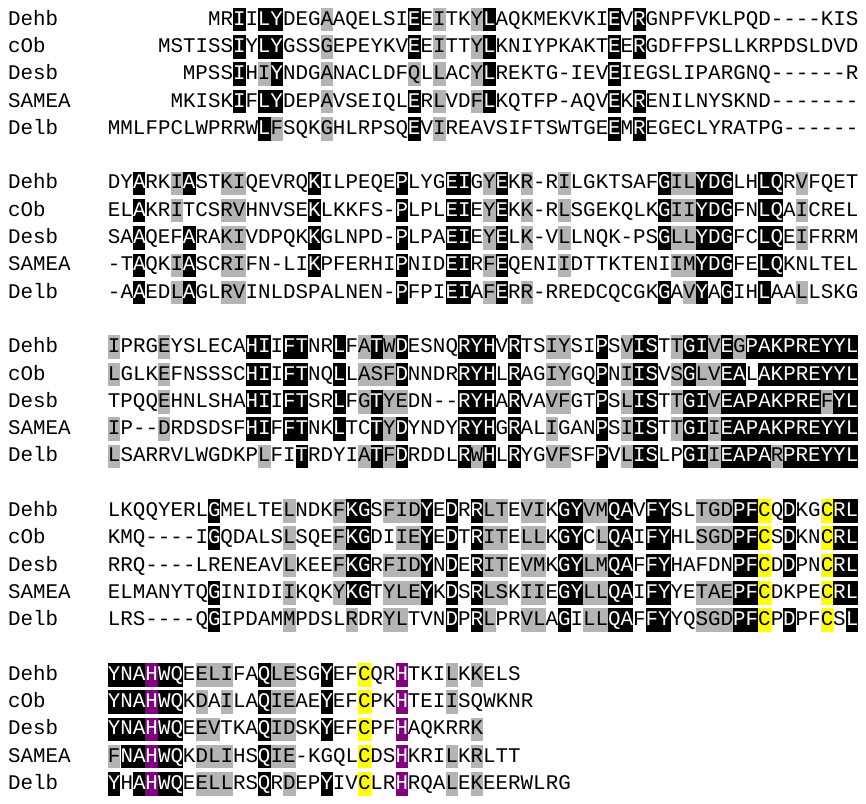

**Figure S3.** Alignment of representative DUF6775 proteins from **Figure 2a** whose genes cluster with suicide Thi4 genes. Potential metal-binding conserved cysteine and histidine residues are highlighted in yellow and purple, respectively. Abbreviations: Dehb, Dehalococcoidales bacterium ARS1330; SAMEA, Archaeal metagenome SAMEA7390951; cOb, *Candidatus* Omnitrophica bacterium isolate bin.255; Delb, Deltaproteobacteria bacterium RBG_16_54_11; Desb, Desantibacteria bacterium DeMMO1_1.

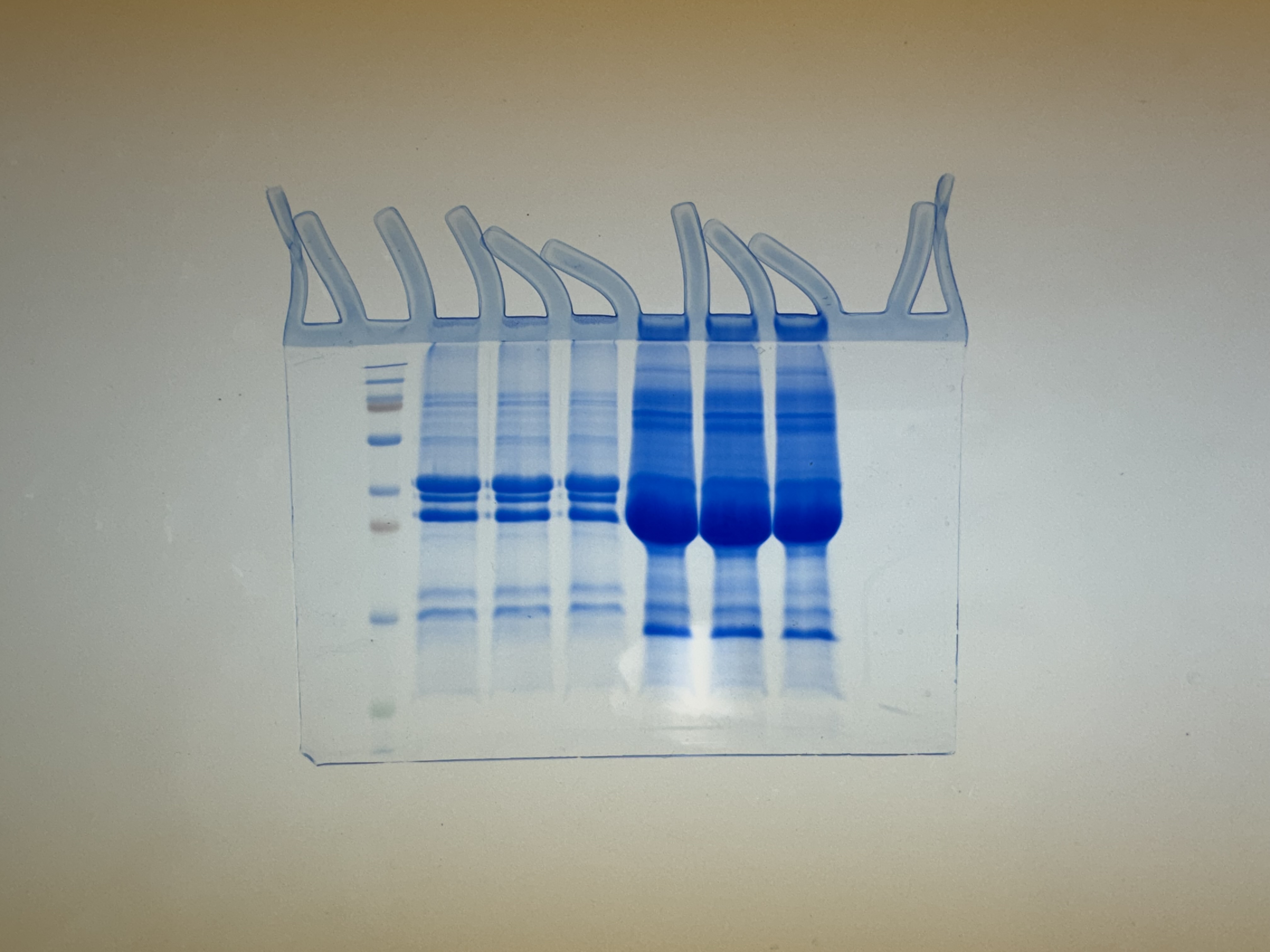

1	2	3

43

M

kDa

Archaeal metagenome SAMEA7390951

34

26

1	2	3

Empty vector

**Figure S4.** SDS-PAGE analysis of recombinant DUF6775 proteins from archaeal metagenome SAMEA7390951. Track labels: M, protein molecular mass markers; EV1-3: empty vector; Arc1-3: triplicates of SAMEA7390951. Staining was with GelCode Blue Safe Protein Stain.

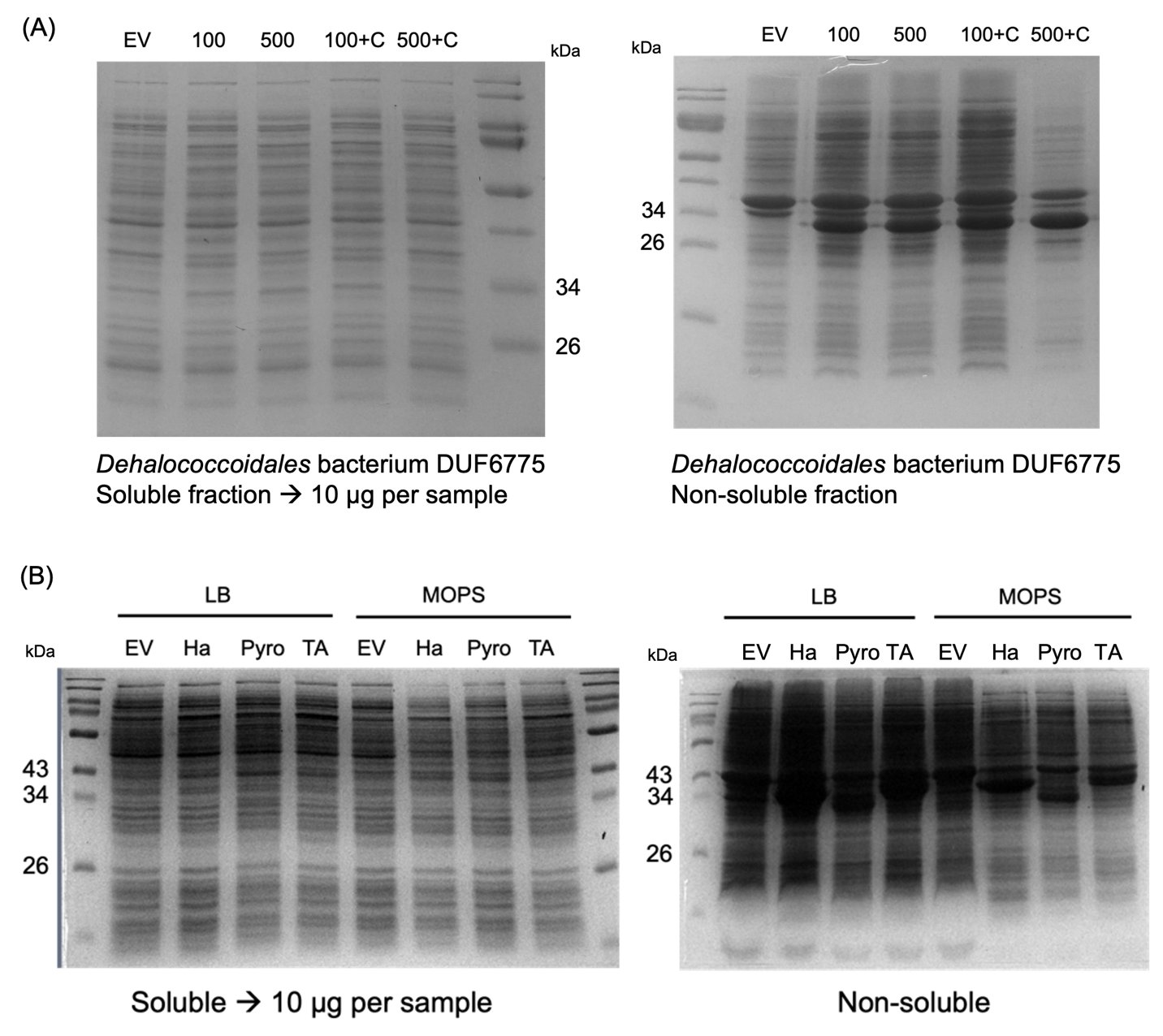

**Figure S5.** SDS-PAGE analysis of recombinant DUF6775 proteins from different sources. **(A)** Expression test of *Dehalococcoidales* bacterium DUF6775 encoding sequence using different IPTG concentrations (100 or 500), with or without the addition of chloramphenicol (indicated by +C). **(B)** Expression tests of different DUF6775 on LB or MOPS media. EV: empty vector; Ha: *Hadesarchaea* archaeon DG-33-1 2656880379; Pyro: *Pyrococcus* sp. NA2 650831530; TA: candidate division TA06 bacterium SM23_40. Staining was with GelCode Blue Safe Protein Stain.
